# Differential binding cell-SELEX method to identify cell-specific aptamers using high-throughput sequencing

**DOI:** 10.1101/466722

**Authors:** Karlis Pleiko, Liga Saulite, Vadims Parfejevs, Karlis Miculis, Egils Vjaters, Una Riekstina

## Abstract

Aptamers have in recent years emerged as a viable alternative to antibodies. High-throughput sequencing (HTS) has revolutionized aptamer research by increasing the number of reads from a few (using Sanger sequencing) to millions (using an HTS approach). Despite the availability and advantages of HTS compared to Sanger sequencing, there are only 50 aptamer HTS sequencing samples available on public databases. HTS data in aptamer research are primarily used to compare sequence enrichment between subsequent selection cycles. This approach does not take full advantage of HTS because the enrichment of sequences during selection can be due to inefficient negative selection when using live cells. Here, we present a differential binding cell-SELEX (systematic evolution of ligands by exponential enrichment) workflow that adapts the *FASTAptamer* toolbox and bioinformatics tool *edgeR*, which are primarily used for functional genomics, to achieve more informative metrics about the selection process. We propose a fast and practical high-throughput aptamer identification method to be used with the cell-SELEX technique to increase the aptamer selection rate against live cells. The feasibility of our approach is demonstrated by performing aptamer selection against a clear cell renal cell carcinoma (ccRCC) RCC-MF cell line using the RC-124 cell line from healthy kidney tissue for negative selection.

## INTRODUCTION

Aptamers are short (20 - 100 nt) oligonucleotides that, contrary to most other functional nucleic acids, bind specific molecular targets due to their folded three-dimensional (3D) structures ^1^. Most aptamers are developed for therapeutic or diagnostic purposes ^2,3^. Several aptamer candidates are currently being tested in clinical trials to treat age-related macular degeneration ^4^, Duchenne muscular dystrophy ^5^, chronic lymphocytic leukaemia ^6^ and other illnesses ^7^.

Aptamer-based diagnostic assays have a great potential to become point-of-care diagnostics: they are affordable, sensitive, specific, user-friendly, robust and can be performed outside a laboratory or hospital. Several commercial aptamer diagnostic platforms have entered the market in recent years ^7^.

OTA-Sense aptamer-based technology to detect ochratoxin, a mycotoxin and potential carcinogen in agricultural products, has been developed by Neoventures Biotechnology Inc. An aptamer-based detection system for aflatoxin is currently marketed as AflaSense. The company is developing similar diagnostic applications for other major mycotoxins including zearalenone, fumonisin and deoxynivalenol ^8^.

ApolloDx’s aptamer-based food pathogen diagnostic platform of food safety testing is marketed by CibusDx. The technology is based on test strips with aptamer-based APOLLOMERTM probes that bind specific targets of foodborne and waterborne pathogens, toxins and viruses present in the test sample ^9^.

The SOMAScan and SOMAmer aptamer array platforms marketed by SomaLogic use modified aptamers with multiplexed proteomics technology enabling high throughput screening of multiple biomarkers in limited sample volumes (150 μl). With SOMAmer technology, more than 1,305 human proteins at less than pg levels in body fluids can be detected ^10^.

The OLIGOBIND©Thrombin activity assay, marketed by Sekisui Diagnostics, GmbH, is a novel oligonucleotide-based enzyme capture methodology that accurately measures thrombin levels through an aptamer-based enzyme-capture fluorescent assay ^11,12^.

AptoCyto, marketed by AptSci, are aptamer products developed for flow cytometry application. Aptocyto technology uses magnetic bead-based cell isolation kits that can efficiently isolate CD-31, EGFR, HGFR, ICAM-2, VEGFR-2 or HER-2 positive cells ^7,13^.

The current aptamer technology market was estimated at USD 1.0 Bn in 2016 with a cumulative annual growth rate of over 20% from 2017 to 2025 ^14^. In comparison, the monoclonal antibodies market size was 85.4 Bn in 2015 and is expected to reach USD 138.6 Bn by 2024 ^15^.

After initially describing an aptamer selection method termed SELEX (systematic evolution of ligands by exponential enrichment) ^16^, several aptamer selection methods have been developed, among others cell-SELEX ^17^, in which live cells are used. The first high-throughput SELEX (HT-SELEX) experiment, a variation of the SELEX process that uses high-throughput sequencing (HTS) methods instead of Sanger sequencing, was described by Zhao et al. in 2009 ^18^. The consequent adaptation of HTS methods for aptamer research further improved selection procedure outcomes ^19,20^.

Subsequently, RNA aptamer selection against the active and inactive conformation of β_2_ adrenoreceptor described by Kahsai et al. employed HTS methods to characterize the fold change enrichment of particular sequences during the selection against each individual target in parallel ^19^. However, in cell-SELEX, this approach might be of very limited use due to the high diversity of protein targets on the cell surface that would cause the enrichment of non-specifically bound sequences if no negative selection were performed.

Several research teams have developed tools to analyse HTS data in aptamer selection, notably *FASTAptamer*, a toolkit developed by Alam et al. that can be used to track the evolutionary trajectory during the SELEX process of individual oligonucleotide sequences ^20^. Recently, *AptaSUITE*, a comprehensive bioinformatics framework that includes most of the previously published functionalities of different tools (data pre-processing, sequence clustering, motif identification and mutation analysis) has been introduced ^21^.

RNA-sequencing (RNA-seq) experiments are used to quantify the differential expression of gene transcripts between samples ^22^. We speculated that it might be possible to adapt data analysis tools currently used for RNA-seq to be used with HT cell-SELEX experiments. During the cell-SELEX experiment, the goal is to select aptamers that bind to the target cells in larger numbers compared to the control cells, making the experimental design similar to RNA-seq analysis. Here, we provide a differential binding cell-SELEX method that can be used to identify differentially abundant aptamers on the surface of target cells and negative control cells during the cell-SELEX experiments and to calculate the statistical significance of these differences. Similar approaches for protein SELEX, in which parallel aptamer selection is performed followed by HTS, have been previously performed to identify broad-spectrum aptamers against the primate lentiviral family of reverse transcriptases ^23^. A comparative binding analysis of a protein SELEX approach in combination with HTS, followed by a detailed characterization of aptamer sequences, has been used to identify high relevance aptamers against serpin plasminogen activator inhibitor-1 ^24^. However, the risk of enrichment of non-specific sequences during cell-SELEX experiments is substantially higher than in protein SELEX due to the inherent complexity of the target in cell-SELEX. While in protein SELEX there is usually only one target, in cell-SELEX the target protein is unknown and selection is performed in live cells with all non-specifically expressed proteins present. Consequently, a statistical analysis of binding differences between the target and negative control cells included in the differential binding cell-SELEX could be a valuable approach for aptamer selection against complex targets. Our analysis includes the use of *edgeR* ^25^, a common tool for analysis in RNA-seq experiments. This tool employs a negative binominal distribution to identify differentially expressed genes. It also employs the *FASTAaptamer* ^20^ toolbox to estimate the read count, *cutadapt* ^26^ to remove the constant primer binding regions of aptamers and a bespoke R script for reuse. Moreover, we combine our approach with a sequence enrichment analysis already used by other groups for aptamer selection to identify the most relevant sequences.

## RESULTS

### Aptamer selection

To identify ccRCC-specific aptamers, the initial randomized oligonucleotide library was subjected to cell-SELEX for 11 selection cycles using the RCC-MF cell line as a target cell line to identify ccRCC-specific aptamers and RC-124 cells as a negative control cell line to reduce the nonspecific binding. Cell-specific aptamer sequence enrichment monitoring was performed using flow cytometry (Guava 8HT) after the 4^th^, 8^th^ and 11^th^ selection cycles. After the 4^th^ and 8^th^ selection cycles, there was a slight difference between the binding of the initial randomized oligonucleotide library compared to the enriched libraries. After the 11^th^ selection cycle, we observed binding of the enriched library to more than >95% of cells. However, the observed binding was nonspecific and the selected aptamer sequences were binding to both the RC-124 (Fig. 1a) and RCC-MF (Fig. 1b) cell lines.

**Figure 1.**
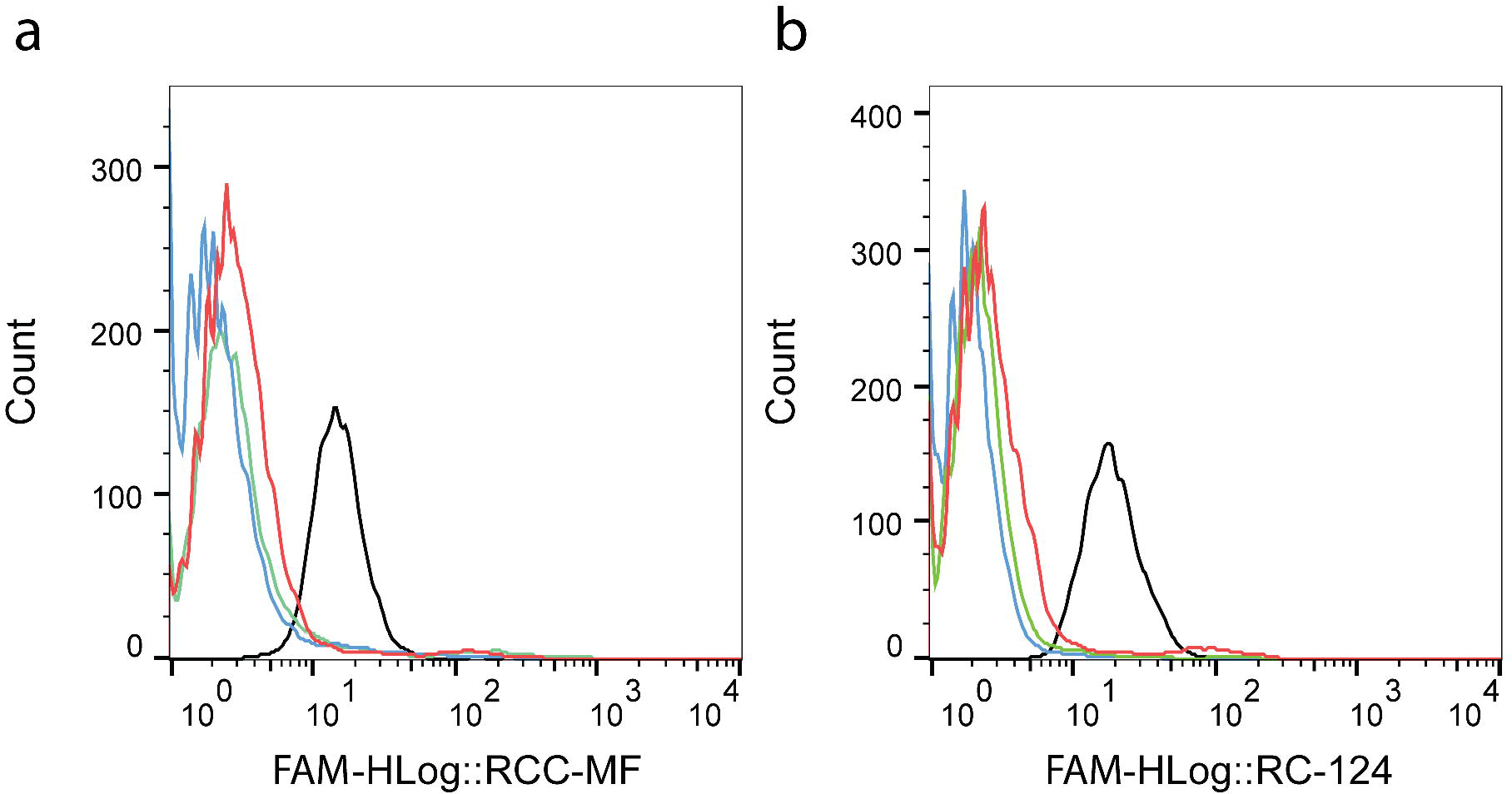
Flow cytometry plots demonstrate fluorescence intensity changes of enriched libraries during the cell-SELEX procedure. Monitoring binding sequences enrichment during cell-SELEX to the negative RC-124 control cells (a) and RCC-MF target cells (b). Blue – randomized oligonucleotide library, green – 4^th^ cycle, red – 8^th^ cycle, black – 11^th^ cycle.

During further selection and process optimization by changing the incubation time, library concentration, FBS concentration and temperature, complete selectivity against the RCC-MF cell line was not achieved up to the 11^th^ cycle.

We concluded that complete selectivity against ccRCC cells is not achieved. However, the low concentration binding measured for the enriched library after the 11^th^ pool (Fig. 2) did not exclude the possibility that the library contains ccRCC cell-specific sequences. To explore the differences that might exist within the library, we developed a differential binding cell-SELEX approach.

**Figure 2.**
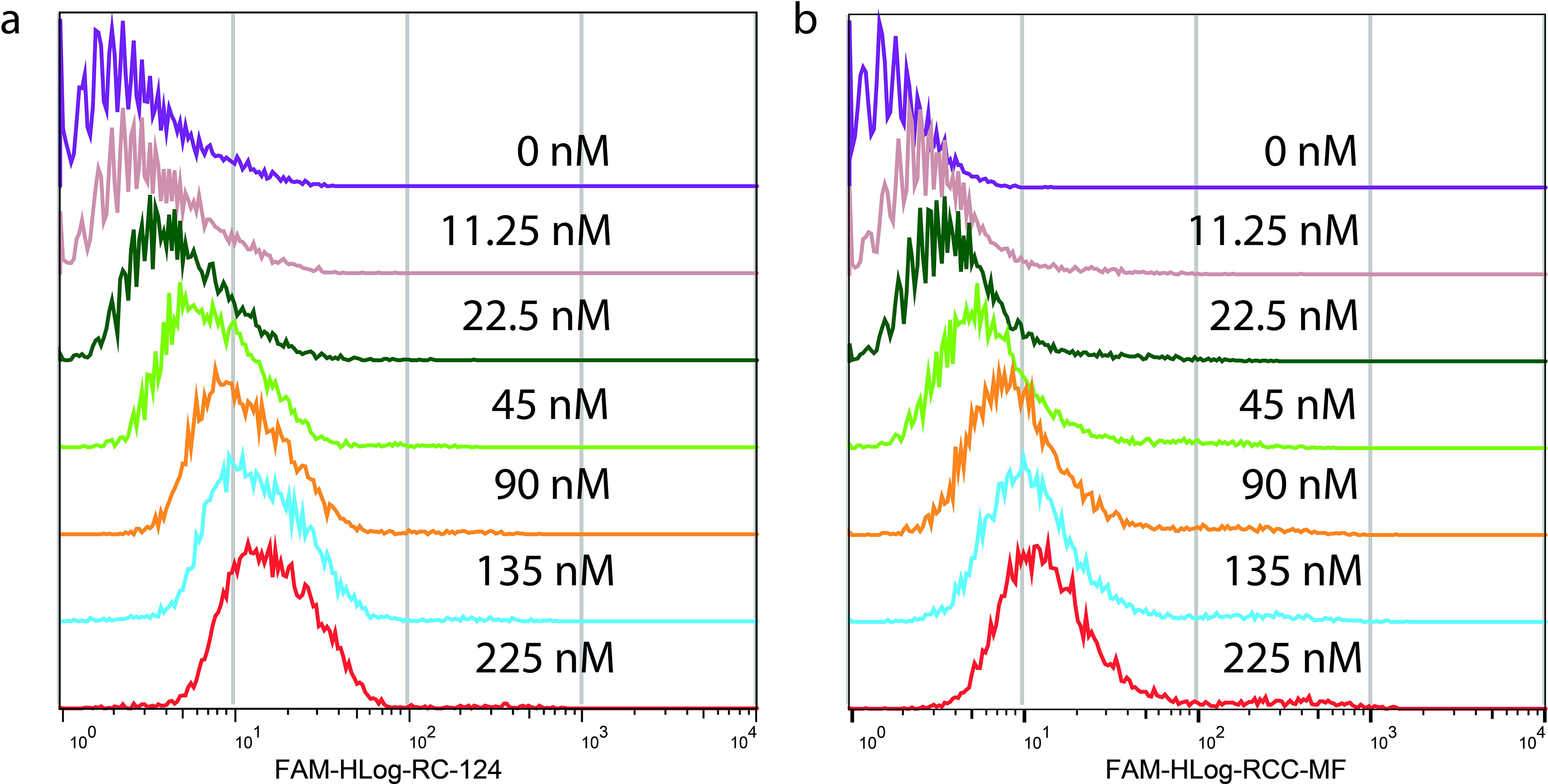
Aptamer binding calculations after the 11^th^ selection cycle. Aptamer binding measurements by flow cytometry using the 11^th^ pool enriched the library at different concentrations on control cells RC-124 (a) and clear cell carcinoma cells RCC-MF (b).

### Differential binding cell-SELEX

The differential binding cell-SELEX process (Fig. 3) was performed after the 4^th^ and 11^th^ selection cycles. After incubation with identically split aptamer libraries and the retrieval of bound sequences to both RC-124 and RCC-MF, we performed two subsequent overlap PCR reactions and confirmed that both constructs after the 1^st^ overhang PCR and 2^nd^ overhang PCR are of expected size (Fig. 4, Fig. S1). Quantification of the final libraries was performed using the NEBNext Library Quant Kit (New England BioLabs) to quantify only those sequences that have flow cell adapters attached to them (Table 1). Overall, our sequencing results also confirm the technical feasibility of the cell-SELEX experiments performed based on the developed protocols (Table 1).

**Table 1.**
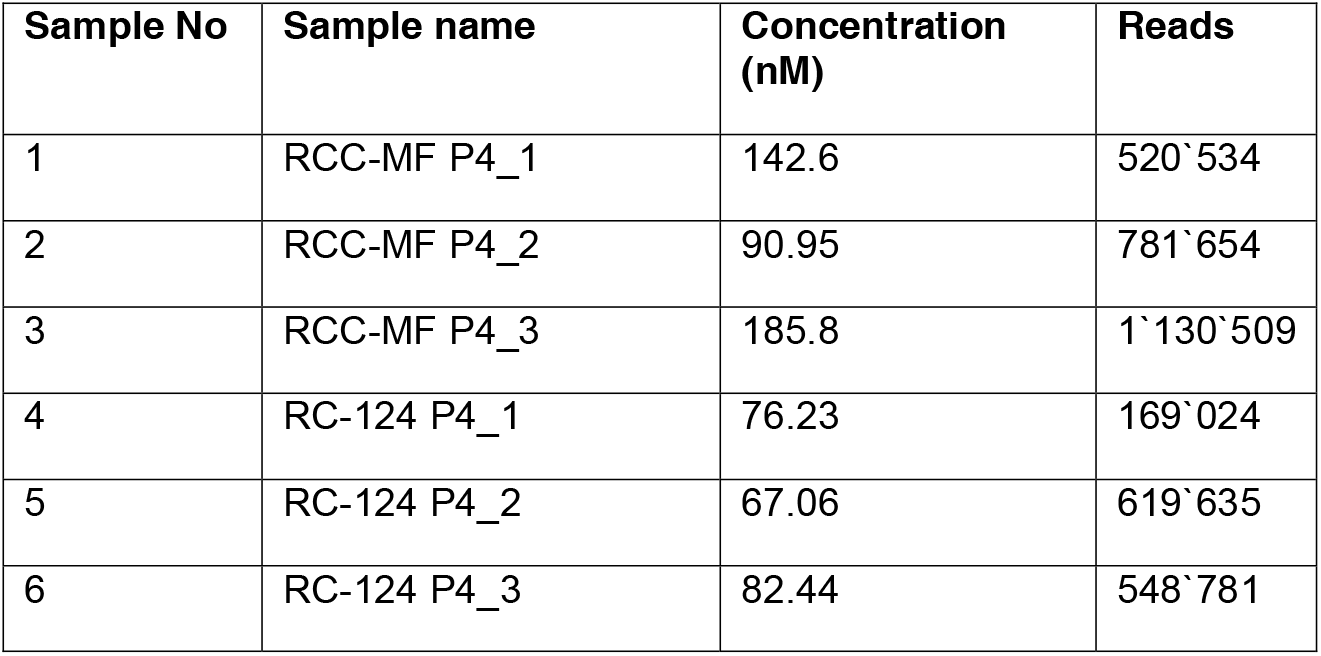
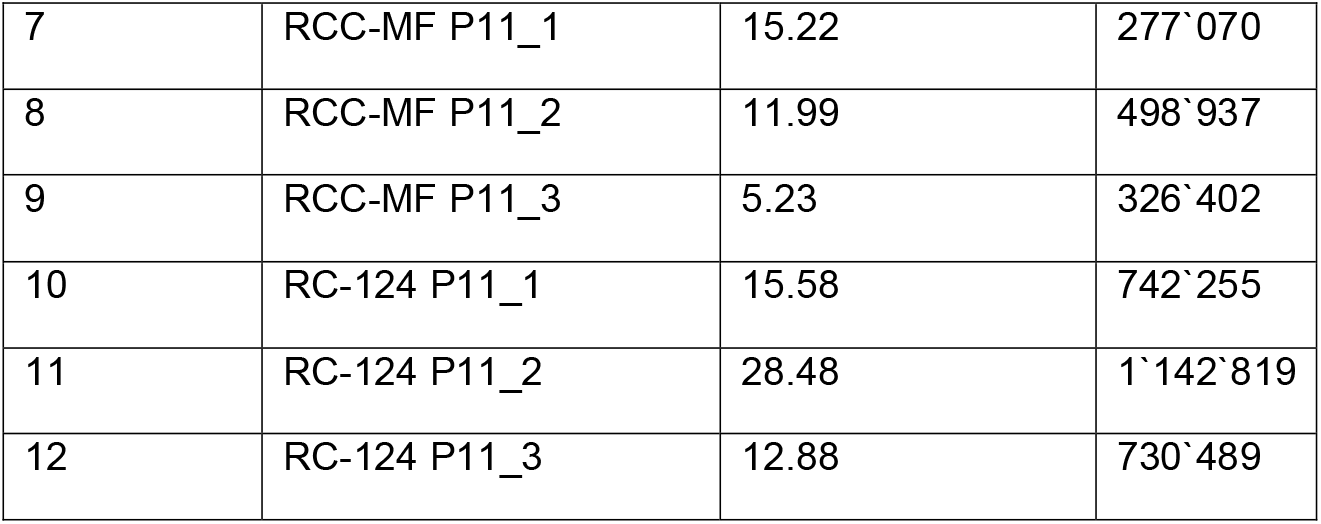
Aptamer concentration determined by qPCR before sequencing and sequencing reads per sample for the sequenced aptamer libraries.

**Figure 3.**
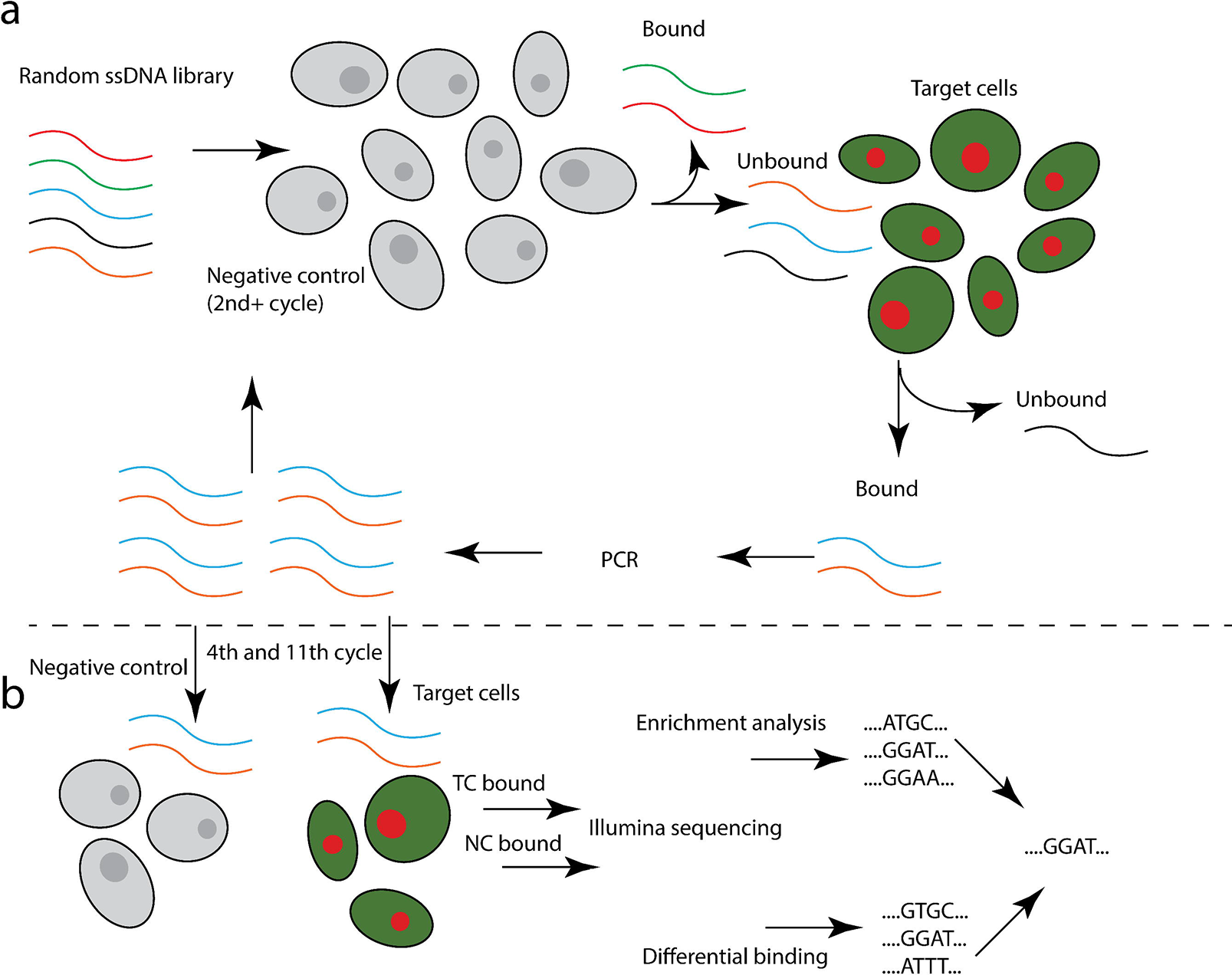
Differential binding cell-SELEX workflow combines (a) the cell-SELEX selection cycle with (b) additional differential binding and data analysis steps to estimate the relative number of aptamer sequences within the pool that bind to each type of cells (TC, target cells; NC, negative control).

**Figure 4.**
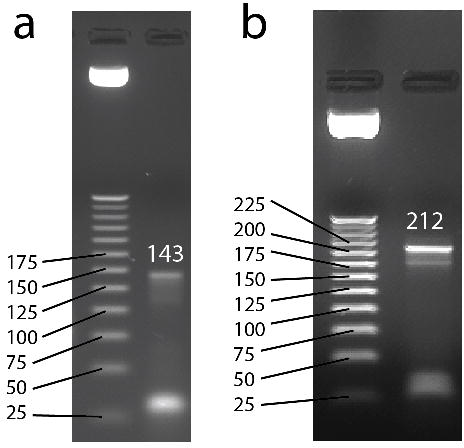
Gel images of aptamers after adding *Illumina* sequencing-specific adapters and indexes. Aptamers after (a) 1^st^ overhang PCR product with a length of 143 bp and (b) 2^nd^ overhang PCR product with a length of 212 bp.

### Data analysis for differential binding cell-SELEX

Sequencing was performed after the 4^th^ and 11^th^ selection cycles. The reads per sample after the initial quality filtration, adapter and constant primer binding region removal and length filtration (40 nt) varied from 169,024 to 1,142,856 (Table 1).

Combining all replicates from both samples after data clean-up, we identified 3,627,938 unique sequences within the 4^th^ selection cycle experiment and 503,107 unique sequences in the 11^th^ selection cycle experiment. After filtering the reads by *edgeR* to remove the sequences that had lower counts per million (CPM) than two per sample and that were present in less than two replicates, we were left with 1,015 unique sequences for the 4^th^ cycle aptamers and 35,859 sequences for the 11^th^ cycle aptamers.

For differential binding data analysis (Fig. 5), we further used selected sequences to run the *edgeR* package, a statistical analysis software that is used to estimate differential expressions from RNA-seq data. The resulting data were adjusted for multiple comparisons using the built-in Benjamini-Hochberg approach and filtered by removing all sequences that have log_2_ fold change (logFC) values less than two or that had adjusted p-values higher than 0.0001.

**Figure 5.**
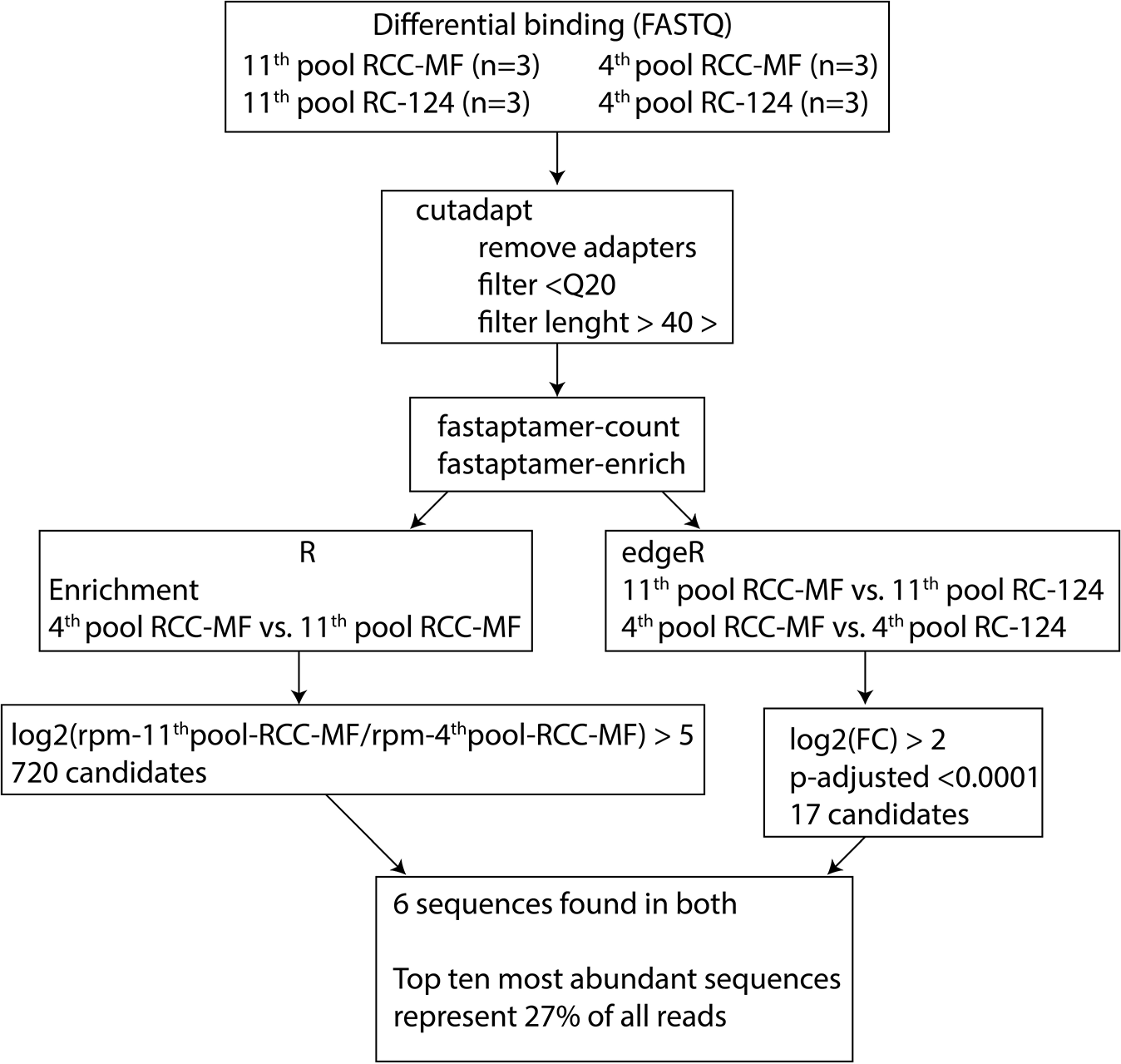
Data analysis pipeline for differential binding cell-SELEX data processing. After trimming using *cutadapt*, the *FASTAptamer* tools *fastaptamer-count* and *fastaptamer-enrich* were used to count the reads for each sequence. Enrichment analysis was performed using the *R* and the *tidyverse* package to identify 720 sequences with enrichment log_2_ > 5. *edgeR* was used to perform differential binding analysis, resulting in 17 candidate sequences. Matching the sequences resulted in six aptamer candidates that are represented in both analyses.

Comparing differential binding datasets using the 4^th^ selection cycle enriched library, we were unable to identify any significantly differentially bound sequences based on the count per million (CPM) of each sequence and a fold change (FC) comparison between two cell lines (Fig. 6a). Most of the sequences bound from the 4^th^ cycle enriched library had a low abundance. However, an analysis of the 11^th^ selection enriched library discovered 195 statistically significant differentially bound sequences according to the same criteria as described for the first experiment (multiple comparison adjusted p-value < 0.0001, log_2_(CPM) > abs(2)) (Fig. 6b). 178 sequences had log_2_(CPM) < −2 compared to 17 sequences that had log_2_(CPM) > 2 (Supplementary Table 1), indicating that more cell type specific sequences were identified for the control RC-124 cells than for the target RCC-MF cells (Fig. 6c).

**Figure 6.**
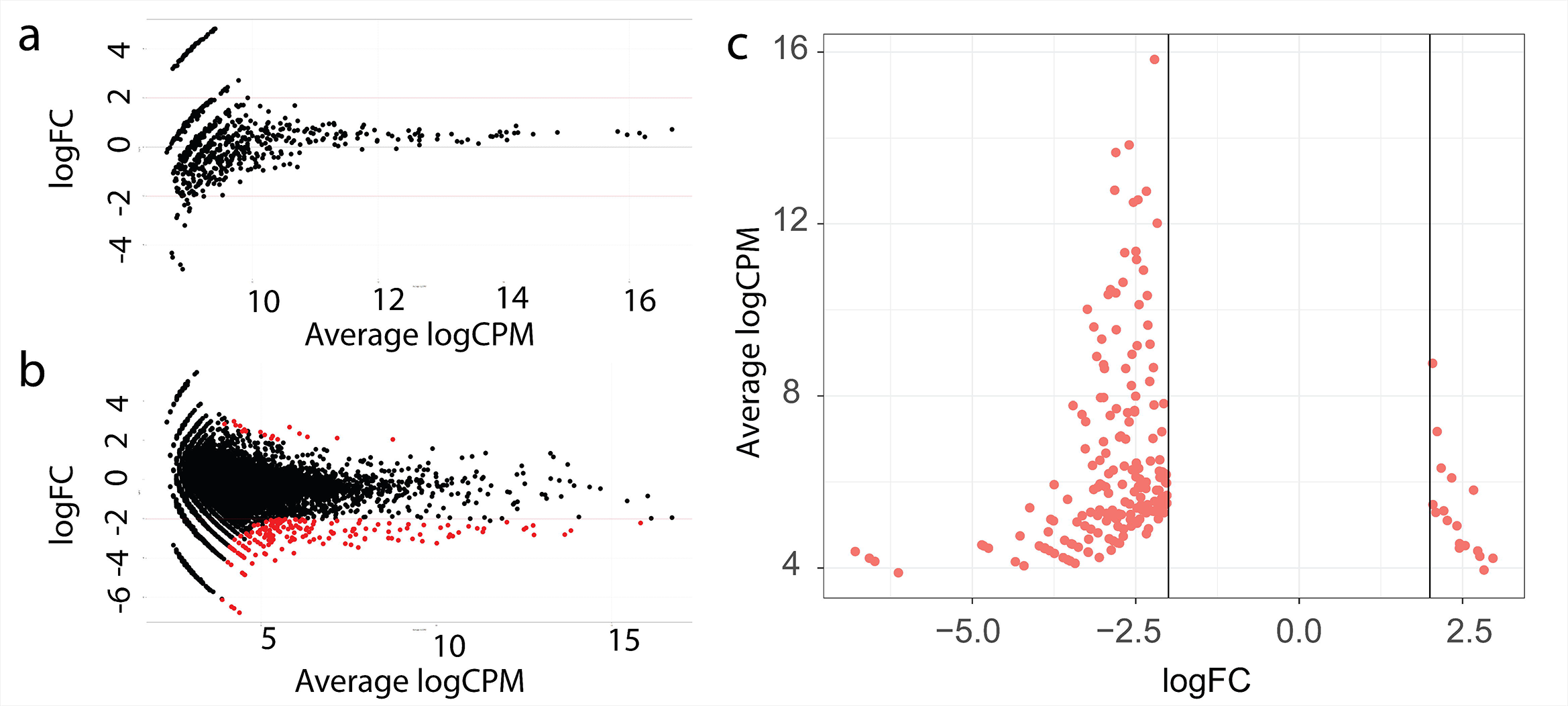
Differential binding cell-SELEX results at the 4^th^ cycle (a) and 11^th^ cycle (b) of selection. A negative logFC value indicates increased binding to the RC-124 control cells, a positive logFC value indicates increased binding to the RCC-MF ccRCC cells, red dots indicate that these results are statistically significant according to an adjusted p-value < 0.0001 using *edgeR* and have logFC > 2 in absolute numbers. All results that fulfil these criteria can be seen in (c).

Enrichment analysis identified 720 unique sequences that have log_2_(meanCPM@11^th^ cycle/meanCPM@4^th^ cycle) > 5 or sequence enrichment in CPM terms 32 times from the 4^th^ to 11^th^ cycle (Supplementary Table 2). We further combined differential binding results that resulted in 17 unique sequences with 720 sequences obtained from enrichment analysis. We identified only 6 sequences that were present in both datasets (Supplementary Table 3) as the most likely candidates to specifically target ccRCC cells (if the log_2_ cut off value is decreased to 5, it is possible to identify 6 sequences that can be found in both the differential binding analysis and enrichment analysis results). We also ordered all unique sequences that were present in the 11^th^ pool by CPM and calculated the log_2_ enrichment value between the 4^th^ and 11^th^ cycle (Supplementary Table 4). Log_2_ enrichment values for the top 10 most abundant sequences ranged from 4.7 to 6.2, and 7 of 10 sequences had a Log_2_ value above 5, meaning that these sequences are also included in the enrichment analysis results. These 10 most abundant sequences contribute to approximately 27% of all sequencing reads from the 11^th^ pool. However, none of the top 10 most abundant sequences passed the statistical significance threshold or FC threshold in the differential binding analysis.

Differential binding results confirm that it is possible to use *edgeR* within our pipeline to identify the most likely candidate molecules for further testing.

### Functional testing of selected lead aptamers

For lead aptamer testing using flow cytometry, we chose 11 sequences identified by different data analysis methods (DB, differential binding; EN, enrichment and MB, most abundant). The top three sequences in each data analysis method were chosen. Differential binding cell-SELEX analysis alone sorted by CPM identified sequences DB-1, DB-2 and DB-3. Differential binding cell-SELEX together with enrichment analysis sorted by log_2_FC identified the DB-3, DB-4 and DB-5 sequences.

Enrichment analysis between the 4^th^ and 11^th^ pools by log_2_CPM enrichment identified sequences EN-1, EN-2 and EN-3. The three most abundant sequences bound to the RCC-MF cells were MB-1, MB-2 and MB-3 (Table 2). We estimated a population shift as a mode of fluorescence intensity (MFI) for each aptamer sample (n=3). The data were corrected by subtracting the MFI from a sample that was incubated with a randomized starting library (MFI_lead-sequence_-MFI_random-library_).

**Table 2.**
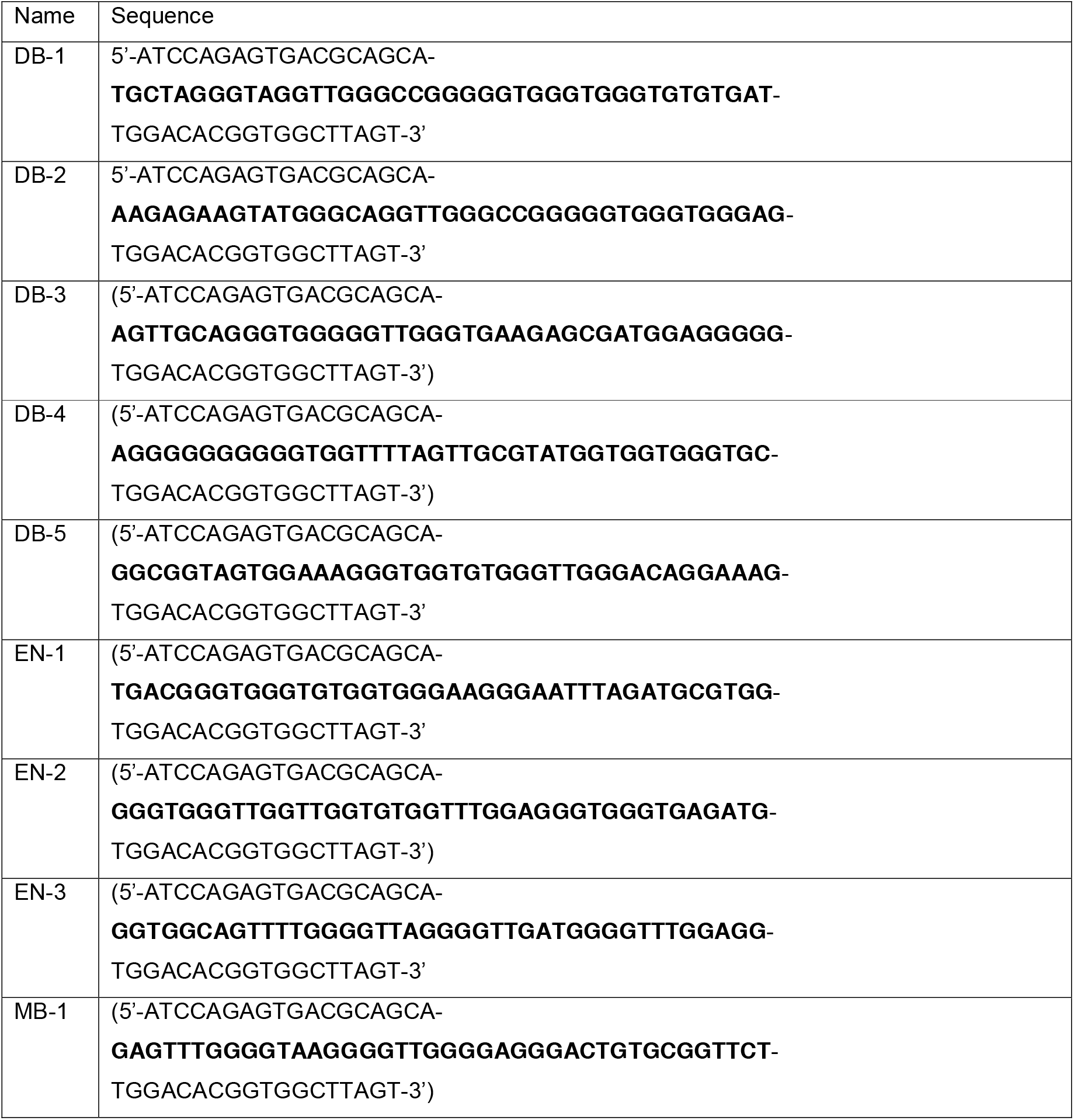
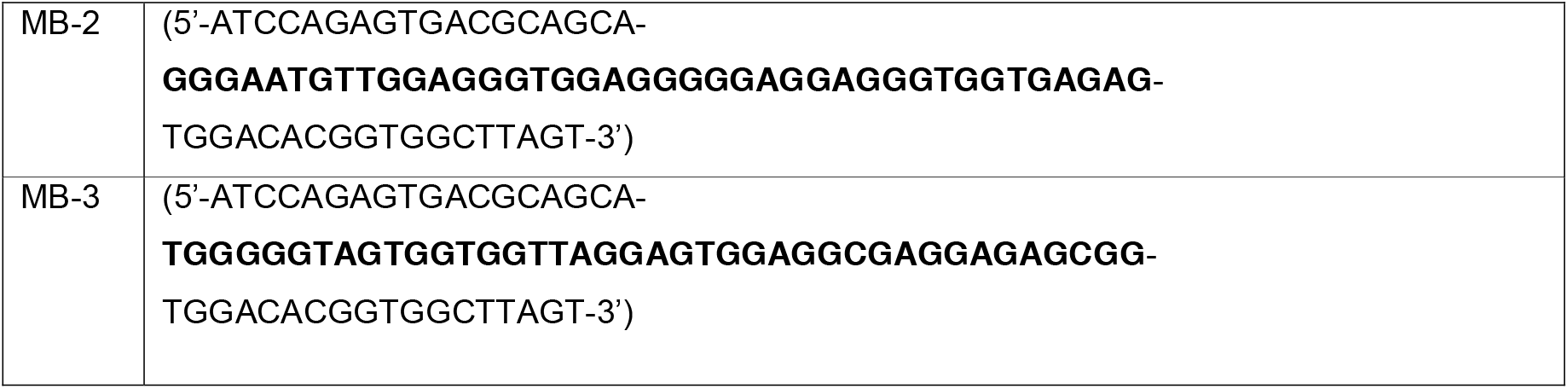
Lead sequences used for the confirmatory cell binding test by flow cytometry.

Corrected MFIs were compared with the t-test (significance defined as p < 0.05, n=3) using GraphPad Prism to determine if our identified sequences altogether bind more to RCC-MF cells than to RC-124 cells. Three sequences (DB-4, EN-2, MB-3) were confirmed to be differentially bound using flow cytometry by comparing MFIs (Fig. 7, Fig. S2). While MB-3, identified as the 3^rd^ most abundant sequence, was significantly (p=0.002) differentially bound, it was targeted towards RC-124 cells. The EN-2 sequence was identified using enrichment analysis and was statistically significantly (p=0.013) binding to RCC-MF cells. DB-4 was significantly (p=0.019) more bound to RCC-MF cells and was identified through a combined differential binding cell-SELEX and enrichment approach.

**Figure 7.**
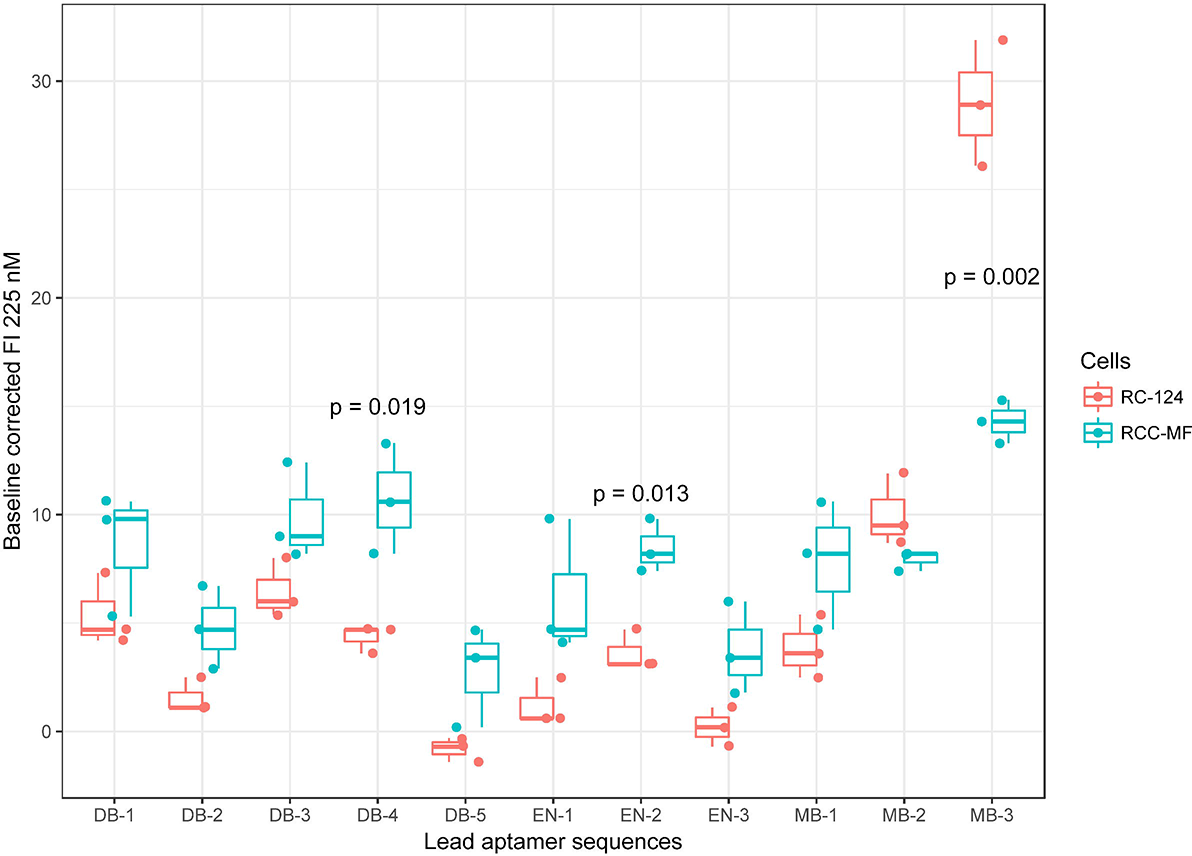
Comparison of the mode of fluorescence intensities from the lead sequences binding to RC-124 and RCC-MF cells. The top three sequences were identified by differential binding alone sorting by CPM (DB-1, DB-2, DB-3), differential binding together with enrichment analysis sorting by log_2_FC (DB-3, DB-4, DB-5), enrichment analysis alone sorting by log_2_CPM enrichment (EN-1, EN-2, EN-3) or by choosing the most abundant sequences in the sequencing dataset (MB-1, MB-2, MB-3). The Upper hinges correspond to the first and third quartiles, and the whiskers mark 1.5* interquartile range (IQR). Statistical significance was determined with a t-test using the GraphPad Prism software.

The binding of selected aptamer sequences was identified through differential binding cell-SELEX (DB-1, DB-2, DB-3, DB-4), and the most abundant sequence (MB-3) was further characterized by flow cytometry analysis. The sequence/randomized library fluorescence intensity ratio at 15 nM, 31 nM, 62 nM, 125 nM, 250 nM, 500 nM and 1000 nM concentrations are plotted in Fig. 8. Less than one ratio was observed for sequences DB-1 and DB-2 when incubated with the RC-124 cell line, indicating that these sequences are binding less than the randomized library to the control cell line. More detailed comparisons for each sequence binding to both cell lines are included in Fig. S3.

**Figure 8.**
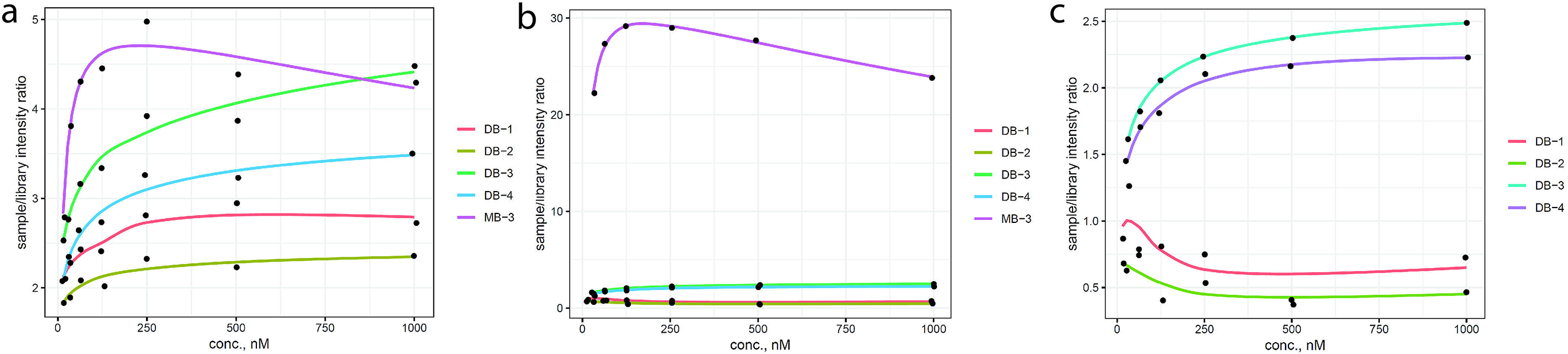
Comparison of fluorescence intensities from flow cytometry for individual sequences (DB-1, DB-2, DB-3, DB-4, MB-3) compared to the randomized library as a ratio (sequence/library). (a) indicates the binding of all sequences to the RCC-MF cell line, (b) includes four sequences (MB-3 excluded for clarity) identified using differential cell-SELEX binding to the RC-124 cell line and (c) characterizes binding to the RC-124 cell line including that of the MB-3 sequence. The y ~ log(x) smoothing function was used when plotting the data.

## DISCUSSION

A recent review on aptamer discovery notes that there are 141 entries of aptamer selection against live cells as of 2017. For comparison, proteins as targets have 584 entries and small molecules have 234 research entries ^1^. This is not surprising considering the advanced technological procedure involved in the cell-SELEX method compared to protein or small molecule SELEX. Several methods have been developed in recent years to improve the success rate of cell-SELEX; for example, HT-SELEX ^18^, FACS-SELEX ^27^ and cell-internalization SELEX ^28^. An HTS adaptation for aptamer sequencing has been described as one of the most fundamental changes to aptamer selection technology ^29^.

The main goal achieved in this research is the development of a differential binding cell-SELEX method. This method can identify cell type-specific aptamer sequences from cell-SELEX selection pools that would not be selected by other cell-SELEX methods and thus would remain overlooked by the investigators.

Currently, the most often used analysis for aptamer finding using HTS data includes enrichment analysis, which means a comparison of the abundance of one particular sequence at the beginning of the SELEX procedure to the abundance of the same sequence after the SELEX procedure.

Enrichment analysis can identify a large number of oligonucleotides with very similar log_2_ enrichment values, as can be seen in our results (Supplementary Table 2). However, enrichment analysis is rarely useful for cell-SELEX because of the high possibility to enrich non-specific sequences. Using enrichment analysis with a cut-off value of log_2_ > 5, we identified 720 sequences to be further tested. However, when the same sequencing dataset was submitted for differential binding analysis using *edgeR*, we identified 17 sequences that were more abundant on the surface of RCC-MF cells than on RC-124 cells.

Enrichment analysis identified one sequence (EN-2) that was significantly (p=0.013) more bound to target RCC-MF cells, as also confirmed by flow cytometry (Fig. 7). We were able to confirm using flow cytometry that another sequence (DB-4), identified with a combined differential binding cell-SELEX and enrichment analysis, was significantly (p=0.019) more bound to the RCC-MF cells. Importantly, DB-4 was found between 720 sequences identified using enrichment analysis, but only as the 528^th^ most enriched sequence. This provides scientific evidence that our approach can be used to identify lead aptamers that most likely would be lost during enrichment analysis.

MB-3, one of the most abundant sequences in the dataset, showed significant binding to RC-124 cells. MB-3 was identified neither in the enrichment analysis results nor in the differential binding cell-SELEX results. However, seven of the 10 most abundant aptamer sequences after the cell-SELEX process were enriched above the set cut-off value log_2_ > 5 and thus did appear in the enrichment analysis results. None of these sequences appeared in the differential binding results because they did not pass the statistical significance test applied to logFC. These observations are in line with previous statements that the most abundant aptamer sequences are not necessarily the best binders ^30^. This proves the value of the differential binding approach for excluding the non-specifically enriched sequences during the cell-SELEX procedure.

After noticing high guanine abundance in several of the identified lead sequences, we searched for G-quadruplex (G4) forming motifs in sequences using QuadBase2 ^31^ TetraplexFinder with high stringency (G_3_L_1-3_) settings. Non-overlapping G4 motifs were identified in three (MB-3, DB-1, DB-2) out of 11 sequences that we previously tested using flow cytometry. Worth noticing is also the fact that the DB-1 and DB-2 sequences were outliers and had below library fluorescence intensity when binding to the negative control RC-124 cell line compared to the other tested sequences. The G4 motifs labelled in *mfold*^32^ predicted relevant aptamer structures (Fig. 9) using 4 °C as a folding temperature, with 5 mM Mg^2+^ and 157 nM Na^+^ concentrations. All sequences contain G4 motifs in randomized regions. Further sequence shortening might be of interest to determine the role of the G4 motifs in these sequences.

**Figure 9.**
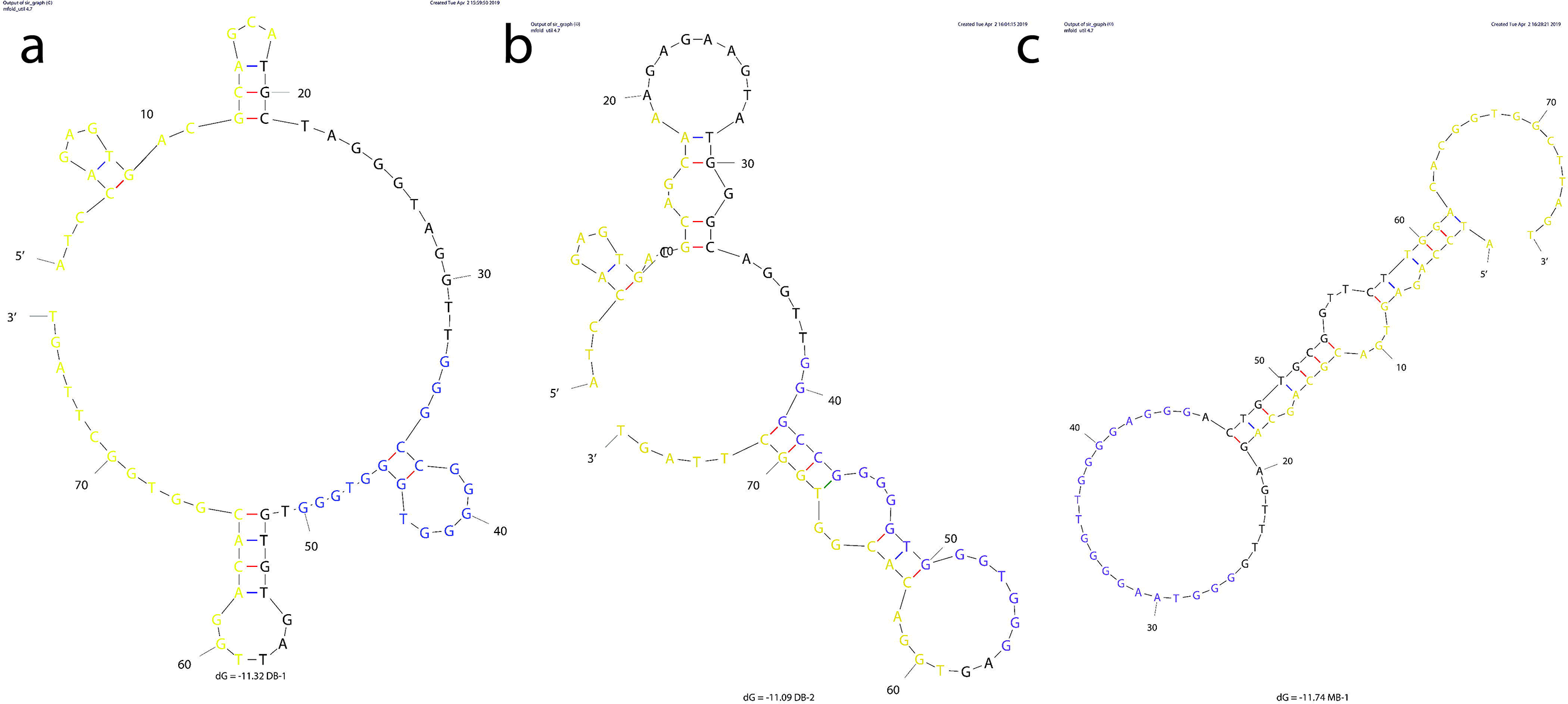
Predicted secondary structures for G-quadruplex containing lead aptamer sequences DB-1 (a), DB-2 (b), and MB-1 (c). Structures were predicted at 4 °C using the mfold web server with 5 mM Mg^2+^ and 157 nM Na^+^ concentrations. Constant sequence regions are highlighted in yellow, and blue represents the identified G4 motifs.

Differential binding cell-SELEX uses *edgeR* to compare how all sequences that can be found in the final enriched aptamer library interact with the control and target cells; it is also used to estimate the statistical significance of these differences. There are several bioinformatics tools available to analyse the statistical significance of the differential expression for RNA-seq data ^25,33,34^. To the best of our knowledge, none of these tools have been applied to estimate differentially bound aptamers on the cell surface. *edgeR* was chosen because it is compatible with the existing data analysis workflows in *R* ^35^. A combination of enrichment analysis and the differential binding approach provides an algorithm to choose target sequences for further analysis.

Altogether, we demonstrate a combined analysis pipeline that can be used to identify lead aptamers from low binding selectivity aptamer libraries after cell-SELEX experiments. We propose a fast and practical high throughput aptamer identification method to be used with the cell-SELEX technique to increase the successful aptamer selection rate against live cells.

A higher number of sequencing reads during differential binding cell-SELEX could even further increase the likelihood to identify low abundance, but differentially bound sequences specific to cells of interest. Sequences that were present only in one replicate from each selection pool were discarded. After the 4^th^ selection cycle, only a few sequences were present in more than one sequencing replicate (Fig. 6a) compared to the 11^th^ cycle (Fig. 6b). An increased number of reads would cover more diverse libraries and would make it possible to identify differentially bound aptamers using fewer selection cycles.

The cell-SELEX design described in this research uses commercially available human RCC-MF and RC-124 cells both as a target and a negative control. We are the first to use these cell lines for aptamer selection with a cell-SELEX approach. However, it could be more suitable to use patient-matched primary cells isolated from the tumour site and adjacent healthy kidney tissue within a few passages after isolation, when cells are most likely to represent the diversity found in clinical settings ^36^.

The differential binding cell-SELEX method developed here can be used to accelerate aptamer selection based on HTS analysis. Additional information from differential binding cell-SELEX reduces the time needed to identify aptamers. This can lead to the broader use of the cell-SELEX technique not only to identify aptamers against cell lines but also against primary cells isolated from patient samples.

We conclude that the differential binding cell-SELEX method can be used to characterize not only sequence enrichment between selection cycles, but also to select aptamer sequences that selectively bind to the target and control cells. We demonstrate the feasibility of our approach by showing cell-line specific aptamer identification against the ccRCC cell line RCC-MF as well as the RC-124 cell line from healthy kidney tissue.

## MATERIAL AND METHODS

### Cell culturing and buffer solutions

Kidney epithelial cell line RC-124 (Cell Lines Service GmbH) established from non-tumour tissue of kidney and carbonic anhydrase 9 (CA9)-positive ccRCC cell line RCC-MF (Cell Lines Service GmbH) established from renal clear cell carcinoma pT2, N1, Mx/ GII-III (lung metastasis) were used for the cell-SELEX process as a negative control and as target cells accordingly. RCC-MF cells were cultured in RPMI 1640 (Gibco), and RC-124 cells were cultured in McCoy’s 5A medium (Sigma-Aldrich). Both culture media were supplemented with 10% foetal bovine serum (FBS) (Gibco), 50 U/ml penicillin and 50 µg/ml streptomycin (Gibco). The cells were propagated at 37 °C, 5% CO_2_ and 95% relative humidity.

Washing buffer containing 4.5 mg/ml D-glucose and 5 mM MgCl_2_ in phosphate-buffered saline (PBS) (SigmaAldrich, D8537, contains K^+^ at 4.45 mM, Na^+^ at 157 mM concentrations) was filtered through a 0.22 µM syringe filter (Corning). The binding buffer contained 4.5 mg/ml D-glucose, 5 mM MgCl_2_, 1 mg/ml bovine serum albumin (SigmaAldrich) and 0.1 mg/ml baker’s yeast tRNA (SigmaAldrich) in phosphate-buffered saline and was filtered through a 0.22 µM syringe filter.

### Oligonucleotide library

A randomised oligonucleotide library with 40 nt and 18 nt constant primer binding regions on both sides of randomized regions (5’-ATCCAGAGTGACGCAGCA-N40-TGGACACGGTGGCTTAGT-3’) was adapted from Sefah et al. ^37^. A FAM label was attached on one primer (5′-FAM-ATCCAGAGTGACGCAGCA-3′) for flow cytometry monitoring, and biotin was attached at the end of the second primer for ssDNA preparation after each cell-SELEX cycle (5′-biotin-ACTAAGCCACCGTGTCCA-3′). Oligonucleotides were ordered from Metabion or Invitrogen.

### Cell-SELEX procedure

Cell-SELEX protocol was adapted from Sefah et al. ^37^. The aptamer library was prepared in binding buffer at a 14 µM concentration for the first selection cycle, heated at 95 °C for 5 min, folded on ice for at least 15 min, and added to fully confluent RCC-MF cells in a 100-mm Petri plate (Sarstedt) that were washed 2 times with washing buffer before the addition of the library. The initial library was applied to RCC-MF cells and incubated for 1 hour on ice with RCC-MF cells but not with RC-124 cells in the first selection cycle. After incubation with the oligonucleotide library, the cells were washed with 3 ml of washing buffer for 3 min and collected with a cell scraper after adding 1 ml of DNase free water. DNase free water was used to collect sequences only for the first cycle; in subsequent cycles, binding buffer was used to retrieve the bound sequences. After collection, the cell suspension was heated at 95 °C for 10 min to remove the bound sequences from the target proteins and centrifuged at 13,000 g; the supernatant containing the selected aptamer sequences was collected.

In subsequent selection cycles, the aptamer library was prepared at a 500 nM concentration and incubated with negative selection cell line RC-124 beforehand. Solution containing unbound sequences was collected and applied to the RCC-MF cell line after washing the cells as described previously. As the selection cycle was increased, a number of modifications were made to the selection procedure: after the 4^th^ selection cycle, 60 mm plates were used instead of 100 mm plates, an increasing concentration of FBS (10-20%) was added to the library after folding without changing the final concentration of the aptamer library, the wash volume was increased to 5 ml, the wash time was increased to 5 min and the number of wash times was increased to 3 after incubation.

### PCR optimization

After each selection cycle, PCR optimization was performed to determine the optimal number of PCR cycles. For PCR optimization and preparative PCR cycling, the conditions involved a 12 min initial activation at 95 °C, followed by repeated denaturation for 30 sec at 95 °C, annealing at 56.3 °C and elongation at 72 °C.

### ssDNA preparation

After preparative PCR, ssDNA was acquired using agarose-streptavidin (GE Healthcare) binding to a biotin-labelled strand, and FAM-labelled ssDNA was eluted with 0.2 M NaOH (Sigma-Aldrich). Desalting was done using NAP5 gravity flow columns (GE Healthcare), the concentration was determined measuring UV absorbance (NanoQuant Plate, M200 Pro, Tecan), and the samples were concentrated using vacuum centrifugation (Eppendorf).

### Monitoring of aptamer binding by flow cytometry

In the enriched aptamer pool, randomized starting library and selected lead aptamers were prepared in binding buffer at 1 µM concentrations, heated to 95 °C for 5 min and then put on ice for at least 15 min. RC-124 and RCC-MF cells were washed with PBS two times and dissociated using Versene solution (Gibco). Then, 50 µL of the enriched aptamer library, starting library, lead aptamers or binding buffer were added to 50 µL of the cell suspension (2.5 * 10^5^ cell per sample), followed by the addition of 11 µL of FBS to each sample to a final concentration of 225 nM. The samples were incubated for 35 min on ice. After incubation, the samples were washed two times with 500 µL of binding buffer and resuspended in 500 µL of binding buffer. The samples were passed through a 40 µM cell strainer before flow cytometry analysis. Flow cytometry data were acquired using a Guava EasyCyte 8HT flow cytometer and analysed using the ExpressPro software (Merck Millipore). Flow cytometry data were analysed using FlowJo software, version 10 (FlowJo). 10,000 gated events were acquired for each sample.

Concentration-dependant binding for sequences DB-1, DB-2, DB-3, DB-4 and MB-3 were performed by preparing each sequence in binding buffer at 2 µM, heating at 95 °C for 5 min and folding on ice for at least 15 min. Subsequent manipulations were performed the same way as for a single concentration monitoring with the exception of preparing variable final concentrations (15 nM, 31 nM, 62 nM, 125 nM, 250 nM, 500 nM, 1000 nM) of each sequence in the cell suspension. Flow cytometry data were acquired using Amnis® ImageStream®XMark II (Luminex). Up to 5,000 single cell gated events were collected for each sample. Data were acquired using the INSPIRE® software and analysed using the IDEAS® software (Luminex).

### Differential binding

Aptamer pools after the 4^th^ and 11^th^ selection cycle were prepared in binding buffer, heated and folded as described for the cell-SELEX procedure at a 1 ml volume with a final concentration of 500 nM. 500 µL were added to both the RC-124 cells and RCC-MF cells grown on 60 mm plates in appropriate cell culture media up to 95% confluence. The aptamer pools were added to the RC-124 and RCC-MF cells and incubated for 30 min on ice, then the cells were washed two times and collected using a cell scraper, heated immediately at 95 °C for 10 min, and centrifuged for 5 min at 13,000 g. The supernatants containing the bound sequences from both cell lines were frozen at −20 °C. Sequencing was done to compare the differential binding profiles of the enriched oligonucleotide libraries obtained from both cell lines.

### Sequencing

The samples for sequencing were prepared by performing two subsequent overlap PCRs as described in the 16S metagenomic sequencing library preparation protocol ^38^. The 1st overlap PCR used primers (5’-TCGTCGGCAGCGTCAGATGTGTATAAGAGACAG-ATCCAGAGTGACGCAGCA-3’ and 5’-GTCTCGTGGGCTCGGAGATGTGTATAAGAGACAG-ACTAAGCCACCGTGTCCA-3’) that are complementary to constant regions of the randomized oligonucleotide library with added overhang that includes an Illumina platform-specific sequence. Conditions for the 1st overlap PCR included 12 min of initial activation, followed by 30 sec at 95 °C, 30 sec at 56.3 °C and 3 min at 72 °C. The cycle number was optimized for each sample to reduce the non-specific amplification. Afterwards, PCR products from one sample were pooled together, concentrated using the DNA Clean & Concentrator (Zymo Research) and run on 3% agarose gel at 110 V for 40 min; the band at 143 bp was cut out and purified using the Zymoclean Gel DNA Recovery kit (Zymo Research).

The second overlap PCR used primers that were partly complementary to the previously added overhang and contained adapters to attach oligonucleotides to the flow cell and i5 and i7 indexes (5’-CAAGCAGAAGACGGCATACGAGAT-[i7 index]-GTCTCGTGGGCTCGG-3’ and 5’-AATGATACGGCGACCACCGAGATCTACAC-[i5 index]-TCGTCGGCAGCGTC-3’). Conditions for the second overhang PCR were 12 min at 95 °C, followed by 5 cycles of denaturation at 98 °C for 10 sec, annealing at 63 °C for 30 sec and elongation at 72 °C for 3 min. After PCR products from one sample were pooled together, the mixture was concentrated using DNA Clean & Concentrator (Zymo Research) and run on 3% agarose gel at 110 V for 45 min; the band at 212 bp was cut out and purified using a Zymoclean Gel DNA Recovery kit (Zymo Research). The concentrations for the final products were determined using the NEBNext Library Quant Kit for Illumina (New England BioLabs) by qPCR.

Sequencing was done on the Illumina MiSeq platform using MiSeq 150-cycle Reagent Kit v3 in single read mode for 150 cycles. 9% of PhiX was added to the run. Sequencing was done at the Estonian Genome Center, Tartu, Estonia.

### Sequencing data analysis

Sequencing reads were filtered and demultiplexed. Constant primer binding regions were removed, and sequences that are longer or shorter than 40 nt were discarded using *cutadapt* ^26^. Counting of recurring sequences was done using *fastaptamer-count*, and matching of the sequences found in replicate samples was done using *fastaptamer-enrich*^20^.

The differential expression analysis tool *edgeR* ^25^ was further used for the analysis of sequencing data. Replicate sequencing samples (n=3) from differential binding cell-SELEX experiments after the 4^th^ and 11^th^ selection cycles were combined, and sequences with low abundance (reads per million < 2 and abundant at all in less than 2 sequencing samples) were filtered out. Normalization was performed based on the reads present in each library. Differential binding was estimated using the *edgeR* function to identify significantly differentially expressed genes using the following parameters: log_2_ fold change (log_2_FC) value > 2, p-value < 0.0001, adjusted for multiple comparisons using the Benjamini & Hochberg ^39^ method.

Enrichment analysis was done separately by using all reads that came from the 4^th^ pool and 11^th^ pool RCC-MF cell binding experiments. We calculated the mean log_2_ value of enrichment (mean counts per million (CPM) for a sequence at the 11^th^ cycle divided by the mean CPM for the same sequence at the 4^th^ cycle) for each sequence and kept the sequences that had log_2_FC > 6 or enrichment between the 4^th^ and 11^th^ cycle.

After these steps, we identified the common sequences in differential binding results and sequence enrichment results to identify the most likely lead aptamer sequences. (*RNotebook* used for 4^th^ cycle differential binding analysis and 11^th^ cycle differential binding analysis, including enrichment analysis, can be found on https://github.com/KarlisPleiko/apta).

## AVAILABILITY

*edgeR* is available as a *Bioconductor* package (http://bioconductor.org/packages/edgeR/), *FASTAptamer* was downloaded from github (https://github.com/FASTAptamer/FASTAptamer), *cutadapt* was installed using *Bioconda* ^40^ RNotebooks for data analysis using *tidyverse* ^41^ are available here: https://github.com/KarlisPleiko/apta

## ACCESSION NUMBERS

Sequencing data are available at SRA under accession number PRJEB28411.

## SUPPLEMENTARY DATA

Supplementary Table 1.

Supplementary Table 2.

Supplementary Table 3.

Supplementary Table 4.

Supplementary Data.

## ACKNOWLEDGEMENT

K.P., U.R., and E.V. conceived and designed the project. K.P., L.S., V.P., and K.M. carried out the experiments. K.P. and U.R. wrote the paper. All read and approved the final manuscript.

