## Supplementary Data for "Differential binding cell-SELEX method to identify cell-specific aptamers using high-throughput sequencing"

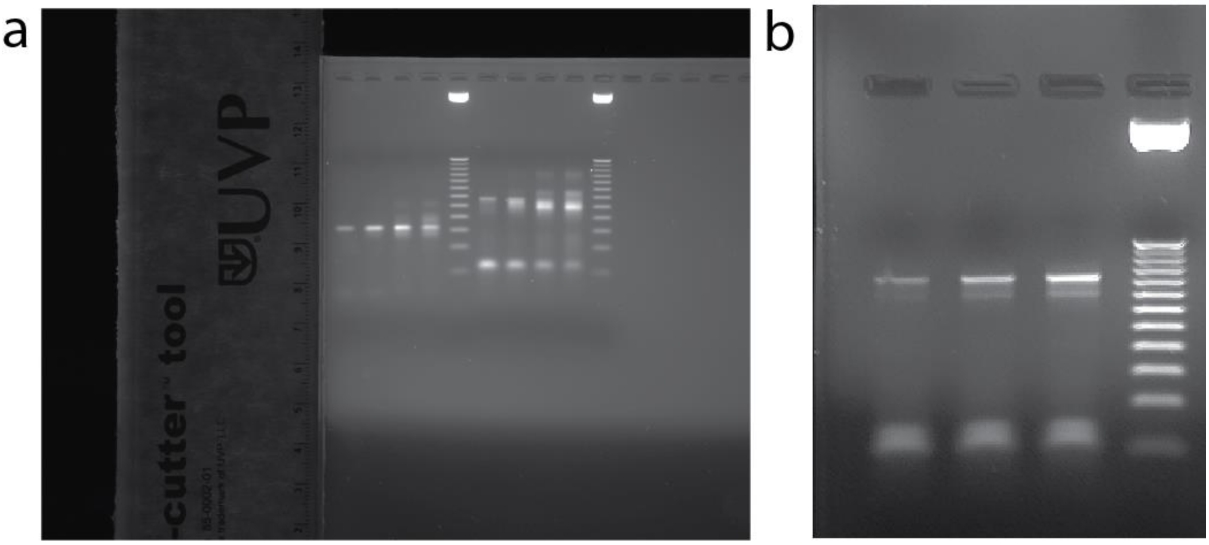


Figure S1. Gel pictures for 1st overhang and 2nd overhang products. a – first four wells from left side contain PCR product amplified using standard primers without overhang, fifth well contains 25 bp size marker, sixth to ninth well contain 1st overhang products amplified using different number of PCR cycles that are comparable to first four wells, 10th well contains 25 bp size marker. Cycle number was chosen based on less byproducts formed (sixth well). b – first three wells contain 2nd overhang products using different concentrations, fourth well contains 25 bp size marker. Cycle number was chosen based on most product observed (third well). Images were acquired using built in UVP software and combined using Adobe Illustrator.

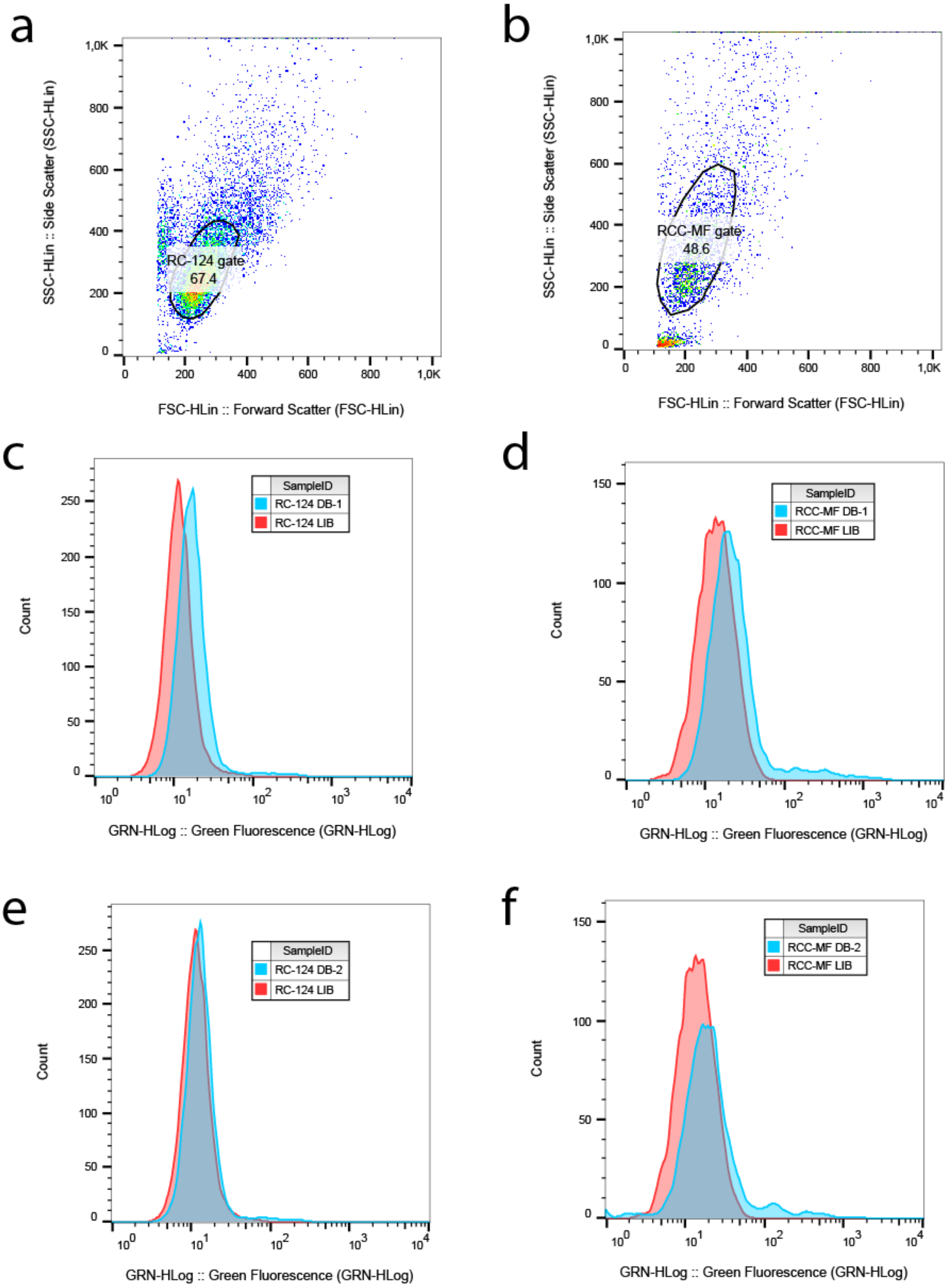
a
SSC-HLin :: Side Scatter (SSC-HLin)
FSC-HLin :: Forward Scatter (FSC-HLin)
RC-124 gate
67.4
b
SSC-HLin :: Side Scatter (SSC-HLin)
FSC-HLin :: Forward Scatter (FSC-HLin)
RCC-MF gate
48.6
c
Count
GRN-HLog :: Green Fluorescence (GRN-HLog)
SampleID
RC-124 DB-1
RC-124 LIB
d
Count
GRN-HLog :: Green Fluorescence (GRN-HLog)
SampleID
RCC-MF DB-1
RCC-MF LIB
e
Count
GRN-HLog :: Green Fluorescence (GRN-HLog)
SampleID
RC-124 DB-2
RC-124 LIB
f
Count
GRN-HLog :: Green Fluorescence (GRN-HLog)
SampleID
RCC-MF DB-2
RCC-MF LIB

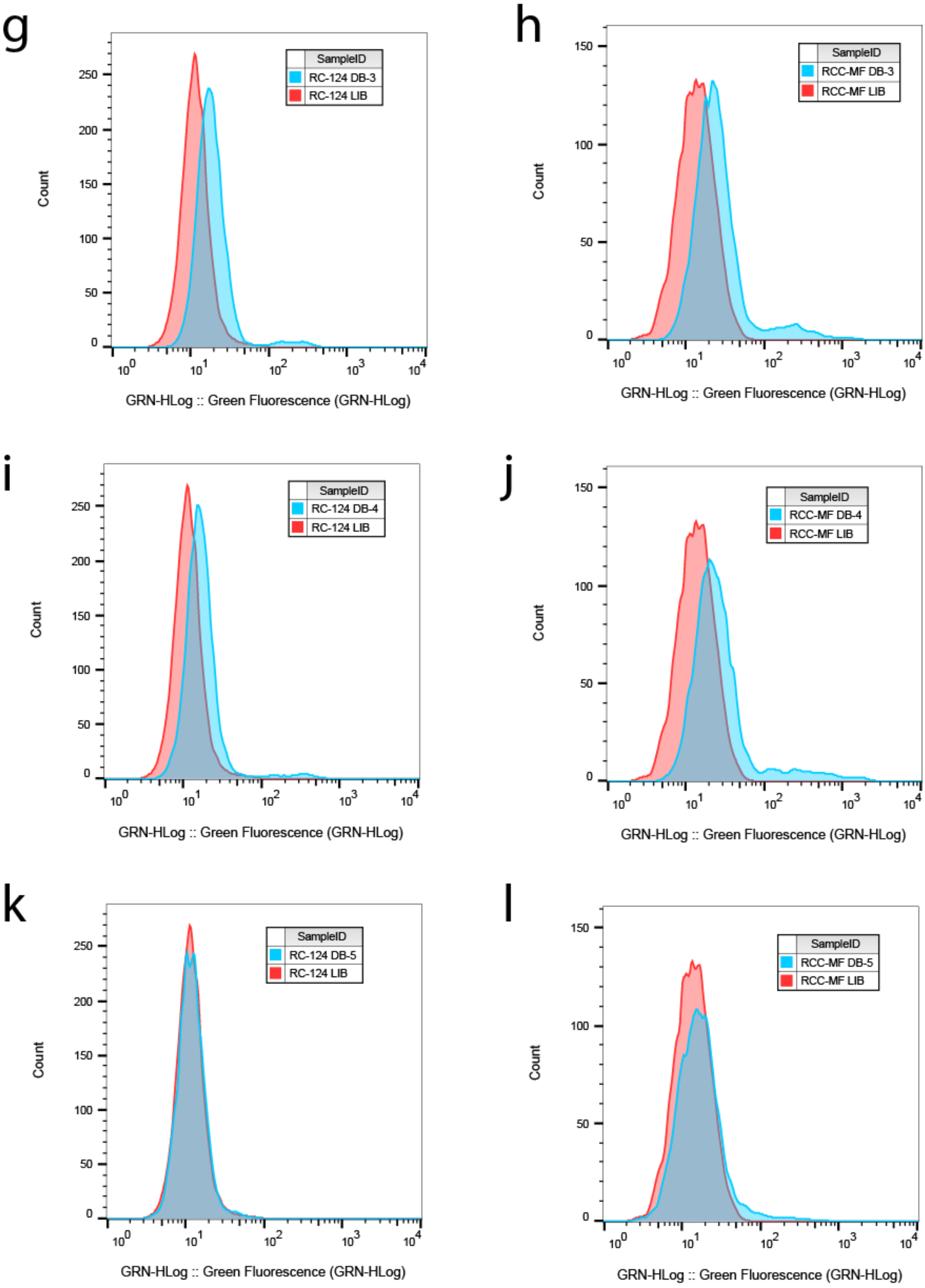
g
SampleID
RC-124 DB-3
RC-124 LIB
Count
GRN-HLog :: Green Fluorescence (GRN-HLog)
h
SampleID
RCC-MF DB-3
RCC-MF LIB
Count
GRN-HLog :: Green Fluorescence (GRN-HLog)
i
SampleID
RC-124 DB-4
RC-124 LIB
Count
GRN-HLog :: Green Fluorescence (GRN-HLog)
j
SampleID
RCC-MF DB-4
RCC-MF LIB
Count
GRN-HLog :: Green Fluorescence (GRN-HLog)
k
SampleID
RC-124 DB-5
RC-124 LIB
Count
GRN-HLog :: Green Fluorescence (GRN-HLog)
l
SampleID
RCC-MF DB-5
RCC-MF LIB
Count
GRN-HLog :: Green Fluorescence (GRN-HLog)

m

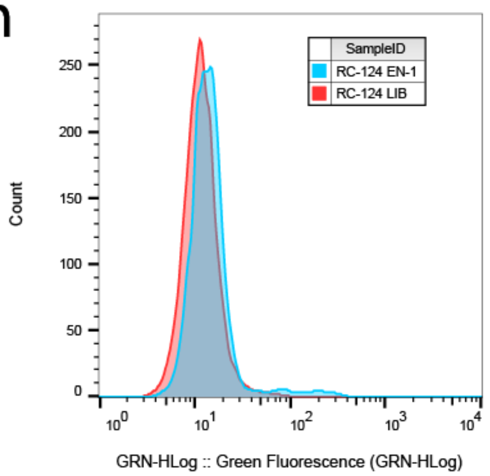

SampleID
RC-124 EN-1
RC-124 LIB
Count
0
50
100
150
200
250
$10^0$
$10^1$
$10^2$
$10^3$
$10^4$
GRN-HLog :: Green Fluorescence (GRN-HLog)


n

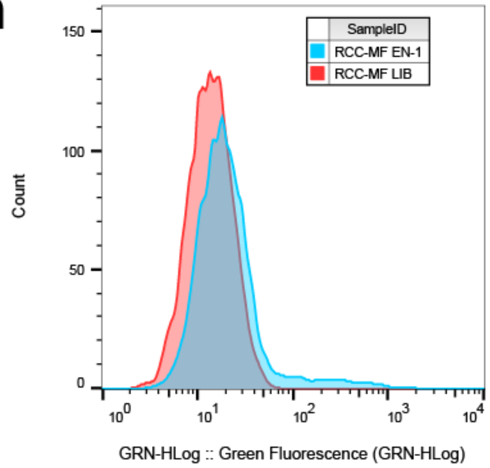

SampleID
RCC-MF EN-1
RCC-MF LIB
Count
0
50
100
150
$10^0$
$10^1$
$10^2$
$10^3$
$10^4$
GRN-HLog :: Green Fluorescence (GRN-HLog)


o

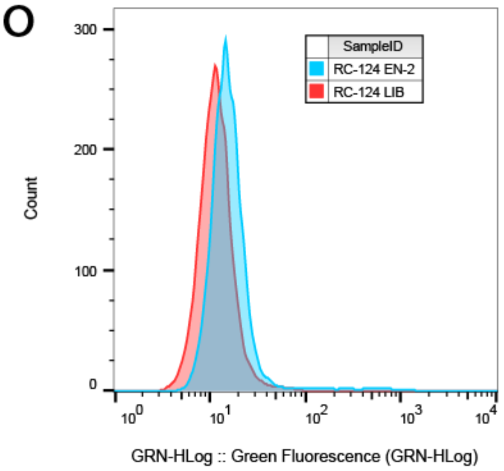

SampleID
RC-124 EN-2
RC-124 LIB
Count
0
100
200
300
$10^0$
$10^1$
$10^2$
$10^3$
$10^4$
GRN-HLog :: Green Fluorescence (GRN-HLog)


p

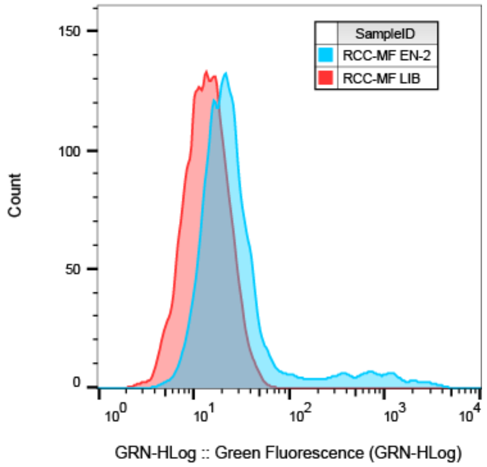

SampleID
RCC-MF EN-2
RCC-MF LIB
Count
0
50
100
150
$10^0$
$10^1$
$10^2$
$10^3$
$10^4$
GRN-HLog :: Green Fluorescence (GRN-HLog)


q

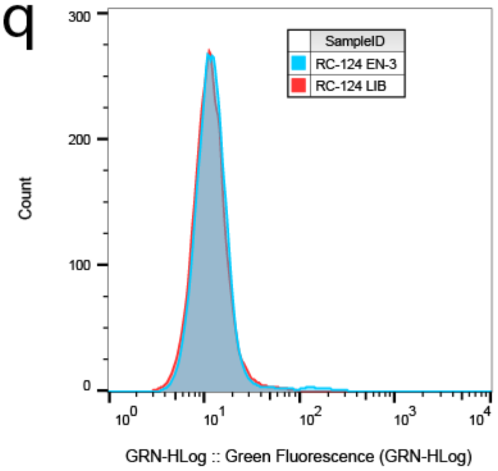

SampleID
RC-124 EN-3
RC-124 LIB
Count
0
100
200
300
$10^0$
$10^1$
$10^2$
$10^3$
$10^4$
GRN-HLog :: Green Fluorescence (GRN-HLog)


r

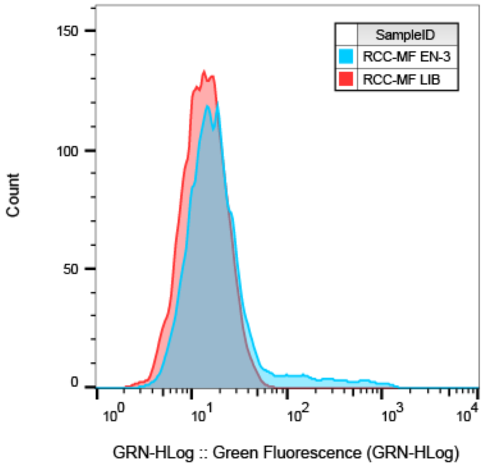

SampleID
RCC-MF EN-3
RCC-MF LIB
Count
0
50
100
150
$10^0$
$10^1$
$10^2$
$10^3$
$10^4$
GRN-HLog :: Green Fluorescence (GRN-HLog)

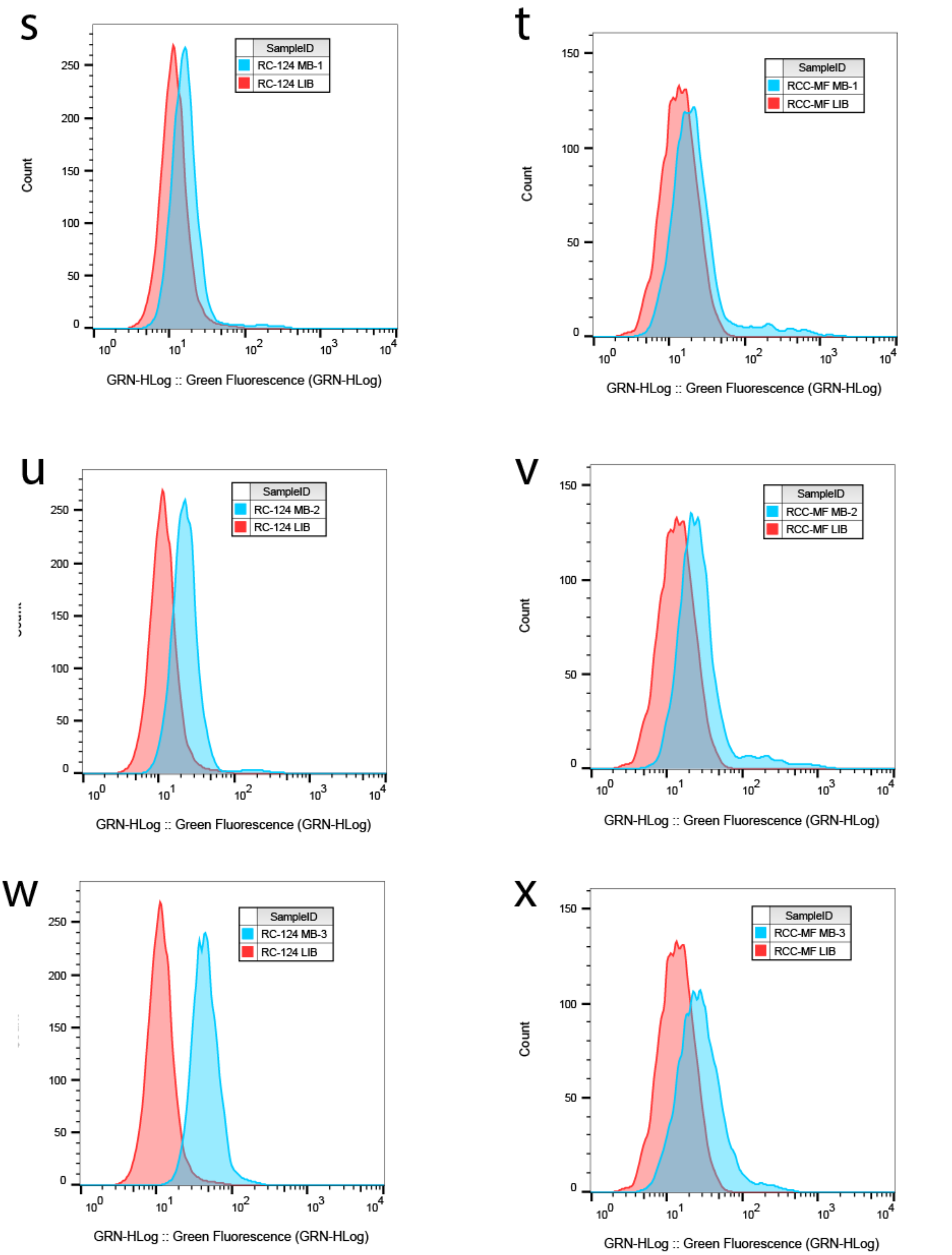


Figure S2. Representative flow cytometry histograms for Figure 7. Gating of RC-124 negative control (a) and RCC-MF target (b) cells. DB-1 sequence binding to RC-124 (c) and RCC-MF (d). DB-2 sequence binding to RC-124 (e) and RCC-MF (f). DB-3 sequence binding to RC-124 (g) and RCC-MF (h). DB-4 sequence binding to RC-124 (i) and RCC-MF (j). DB-5 sequence binding to RC-124 (k) and RCC-MF (l). EN-1 sequence binding to RC-124 (m) and RCC-MF (n). EN-2 sequence binding to RC-124 (o) and RCC-MF (p). EN-3 sequence binding to RC-124 (q) and RCC-MF (r). MB-1 sequence binding to RC-124 (s) and RCC-MF (t). MB-2 sequence binding to RC-124 (u) and RCC-MF (v). MB-3 sequence binding to RC-124 (w) and RCC-MF (x).

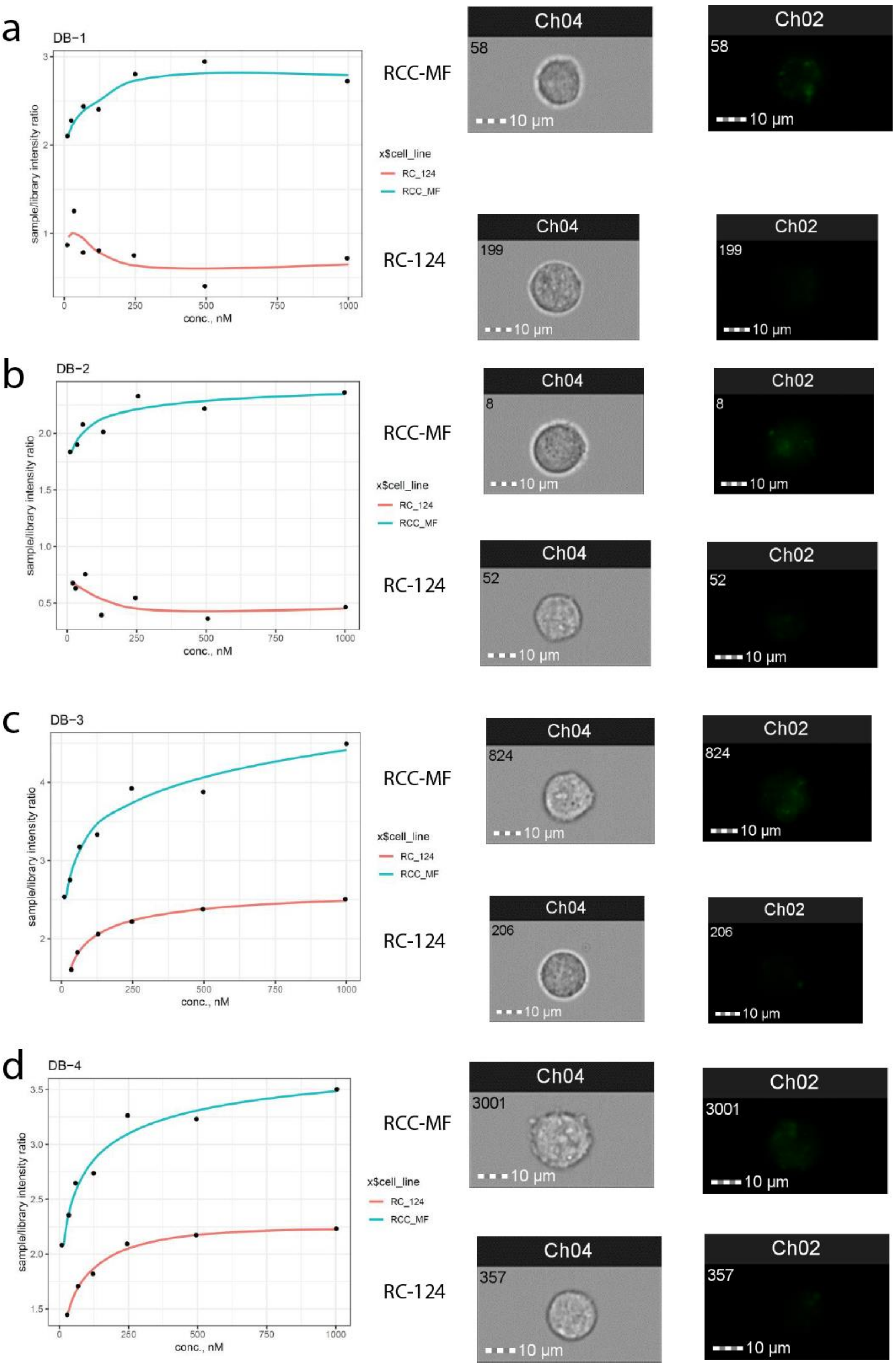
a
DB−1
sample/library intensity ratio
conc., nM
x$cell_line
RC_124
RCC_MF
RCC-MF
RC-124
Ch04
Ch02
58
199
10 µm
b
DB−2
sample/library intensity ratio
conc., nM
x$cell_line
RC_124
RCC_MF
RCC-MF
RC-124
Ch04
Ch02
8
52
10 µm
c
DB−3
sample/library intensity ratio
conc., nM
x$cell_line
RC_124
RCC_MF
RCC-MF
RC-124
Ch04
Ch02
824
206
10 µm
d
DB−4
sample/library intensity ratio
conc., nM
x$cell_line
RC_124
RCC_MF
RCC-MF
RC-124
Ch04
Ch02
3001
357
10 µm

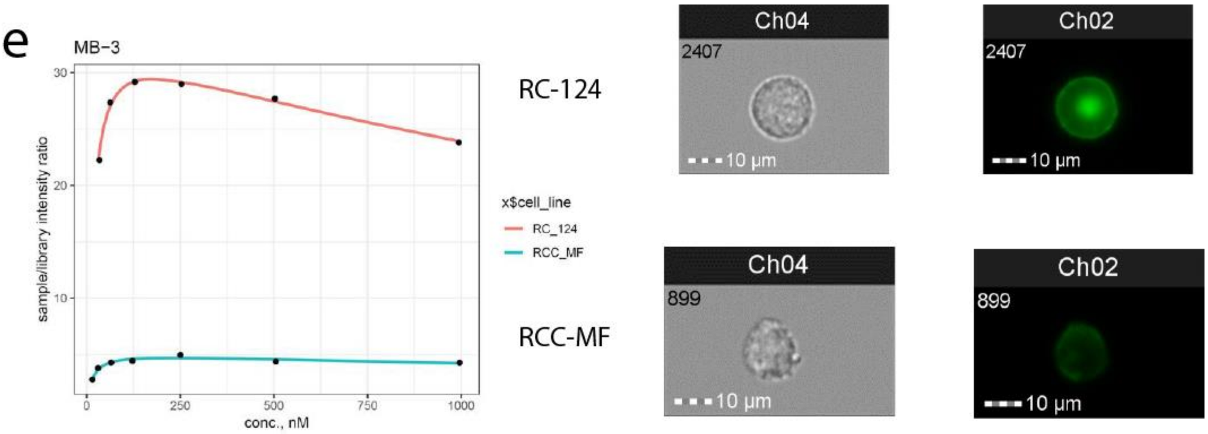


Figure S3. Detailed comparison for data in Figure 8. Concentration dependent binding of sequences DB-1 (a), DB-2 (b), DB-3 (c), DB-4 (d), MB-3 (e) to RCC-MF target and RC-124 negative control cells.
